# Genome sequence of *Pseudomonas aeruginosa* PAO1161, a PAO1 derivative with the ICE*Pae*1161 integrative and conjugative element

**DOI:** 10.1101/494302

**Authors:** Adam Kawalek, Karolina Kotecka, Magdalena Modrzejewska, Grazyna Jagura-Burdzy, Aneta Agnieszka Bartosik

## Abstract

*Pseudomonas aeruginosa* is a cause of nosocomial infections, especially in patients with cystic fibrosis and burn wounds. PAO1 strain and its derivatives are widely used to study the biology of this bacterium, however recent studies demonstrated differences in the genomes and phenotypes of derivatives from different laboratories.

Here we report the genome of *P. aeruginosa* PAO1161 laboratory strain, a *leu*-, Rif^R^, restriction-modification defective PAO1 derivative, described as the host of IncP-8 plasmid FP2, conferring the resistance to mercury. Comparison of PAO1161 genome with PAO1-UW sequence revealed an inversion of a large genome segment between rRNA operons and 100 nucleotide polymorphisms, short insertions and deletions. These included a change in *leuA*, resulting in E108K substitution, which caused leucine auxotrophy and a mutation in *rpoB*, enhancing rifampicin resistance. Nonsense mutations were detected in *PA2735* and *PA1939* encoding a DNA methyltransferase and a putative OLD family endonuclease, respectively. Moreover, a 12 kb RPG42 prophage and a novel 108 kb PAPI-1 like integrative conjugative element (ICE) encompassing a mercury resistance operon were identified. The ICE*Pae*1161 was transferred to *Pseudomonas putida* cells, where it integrated in the genome and conferred the mercury resistance. This indicates that the FP2 plasmid is in fact an ICE.

## INTRODUCTION

*Pseudomonas aeruginosa* is a Gram-negative gammaproteobacterium commonly found in various ecological niches and characterized by the ability to survive in unfavourable, frequently changing environmental conditions. This opportunistic human pathogen is often a source of nosocomial infections in immuno-compromised patients. In cystic fibrosis patients *P. aeruginosa* chronically colonizes the lungs and is a major factor causing mortality^1,2^.

Majority of research on this metabolically versatile bacterium uses sublines or derivatives of *P*. *aeruginosa* PAO1 strain, originally isolated from a wound of a patient in Holloway’s laboratory, Melbourne, Australia^3^. Over the years, the strain was shipped to laboratories worldwide and its different attenuated derivatives, including auxotrophic strains and strains with mobile genetic elements were obtained^4^. In 1999 the genome of *P. aeruginosa* PAO1 stored at the University of Washington (PAO1-UW) was sequenced^5^, providing a reference for studies on *P. aeruginosa* genomes. Up to December 2018, the Pseudomonas Genome Database, a database devoted to the information on *Pseudomonas* species^6^, contained 2226 sequenced *P*. *aeruginosa* genomes, including 11 PAO1 derivatives (sublines). Remarkably, sequencing of the PAO1 subline (MPAO1) as well as PAO1-DSM strain stored at the German Collection for Microorganisms and Cell Cultures revealed presence of multiple nucleotide polymorphisms and short insertions-deletions relative to the reference PAO1-UW^7^. A major feature differing genomes of PAO1 derivatives is the occurrence of a large inversion resulting from the homologous recombination between two rRNA operons *rrnA* and *rrnB*^5^, which is present in the reference PAO1-UW genome and not in e.g., MPAO1 and PAO1-DSM^7^. Despite an asymmetrical positioning of the *dif* region in PAO1-UW, this inversion does not seem to affect chromosome segregation and such large rearrangements might be common among bacteria^8^. The presence of multiple short sequence modifications could lead to the variations in e.g. virulence and fitness between strains used in different laboratories^7^. This data indicate an ongoing micro- and macro- evolution of bacterial genomes and suggests that sequence diversification in laboratory strains should be taken into consideration in the analysis of phenotypic data^9,10^.

In this work we focus on the genome of *P. aeruginosa* PAO1161 strain, a PAO1 derivative requiring leucine for growth on minimal media and selected as lacking the restriction-modification system (*rmo-*10 mutation)^11^. This strain is described as the host for plasmid FP2 of IncP-8, the only known member of this incompatibility group, conferring resistance to mercury^12,13^. The FP2 factor demonstrated the chromosome-mobilizing ability (Cma) and was extensively used in interrupted mating technique for preparation of the genetic map of *P. aeruginosa* chromosome^4,14^. The PAO1161 derives from the PAO38 *leu*-*38* mutant obtained by treatment of PAO1 with manganese chloride and search for leucine auxotrophs^3^ (Figure 1A). PAO38 acquired the FP2 element from PAT (*P. aeruginosa* strain 2)^15^ to yield strain PAO170^16^. Following mutagenesis of PAO170 with N-methyl-N’-nitro-N-nitrosoguanidine PAO1161 was selected as defective in restriction and modification systems (*r*^*-*^*m*) on the basis of the altered susceptibility to phage infection^11,17^. To facilitate the use of PAO1161 in conjugation experiments spontaneous rifampicin resistant clone was obtained^18^. The PAO1161 strain was used in studies on chromosome segregation and gene expression in *P. aeruginosa* using genome wide approaches^19–21^ as well as in other physiological and genetic studies^22–28^.

**Figure 1.**
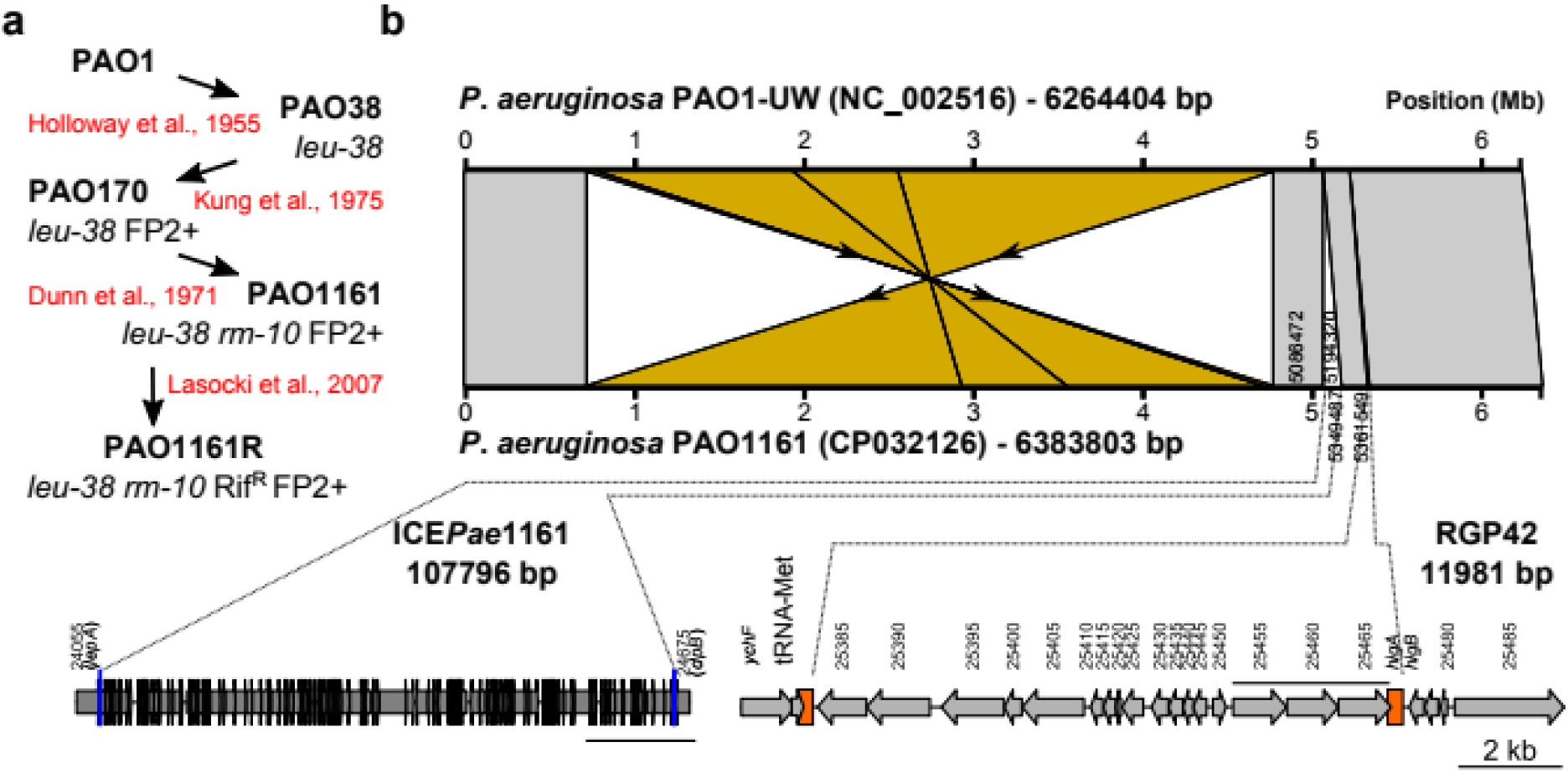
Comparison of *P. aeruginosa* PAO1161 and PAO1-UW genomes. **(A)** Origin of *P. aeruginosa* PAO1161 strain. **(B)** Major structural variations between the genomes of the two *P. aeruginosa* strains. Whole genome alignment and synteny visualization was performed with EasyFig^75^. Blocks indicate regions with percentage of nucleotide sequence identity higher than 95%. The inversion between *rrnA* and *rrnB* rRNA operons is coloured in yellow. Bottom panel indicates positions and schematic gene organization of large insertions: ICE*Pae*1161 and RGP42.

Here we report the genome sequence of *P. aeruginosa* PAO1161 strain. Comparison with PAO1-UW reference sequence revealed the presence of a large number of single-nucleotide polymorphisms (SNPs), insertions-deletions (indels), as well as inversion of large genome segment between rRNA genes. Moreover a functional PAPI-1 like integrative conjugative element (ICE), containing a mercury resistance operon was identified in PAO1161 genome, indicating that the FP2 is not a plasmid but an ICE (designated ICE*Pae*1161).

## RESULTS AND DISCUSSION

### Comparison of *P. aeruginosa* PAO1161 genome with PAO1 reference assembly

*P. aeruginosa* PAO1161 genome was sequenced and assembled, resulting in a single circular chromosome of 6383803 bp. A comparison with the reference PAO1-UW genome (NC_002516) revealed three major structural differences (Figure 1B). The PAO1161 genome lacks the large inversion between ribosomal RNA operons *rrnA* and *rrnB* similarly to other PAO1 derivatives like MPAO1 and PAO1-DSM^7^, but observed in PAO1-UW^5^. Remarkably, PAO1161 possesses two large insertions. The 107796 bp insertion in *tRNA*_Lys_ gene between *lepA* (*PA4541*) and *clpB* (*PA4542*), flanked by 48 bp repeated sequences, displays a significant similarity to PAPI-1 like integrative conjugative elements (see below)^29,30^. The second 11981 bp insertion between *tRNA*_Met_ (*PA4673.1*) and *higA* (*PA4674*), flanked by 82 bp repeats, is identical to the prophage-like RGP42 element identified also in MPAO1 and PAO1-DSM^7^. Additionally, PAO1161 lacks a 280 bp fragment containing *PA1796.3* and *PA1796.4* tRNA genes and has an 107 bp insertion downstream of *PA2327*.

A comparison of PAO1161 and PAO1-UW genome sequences using Nucdiff^31^ revealed 251 single nucleotide polymorphisms (SNPs), multiple nucleotide polymorphisms (MNPs) and short insertions / deletions (indels). Since a direct sequence comparison does not provide information about the quality of predicted variants, the outcome can be greatly affected by errors during genome sequence consensus calling, caused for instance by mapping of reads derived from highly similar sequences to another, very similar, parts of the genome. To identify high quality variants we aligned the reads used in genome assembly to the consensus sequence and checked the frequency of the variant at a given genome position (see Materials and Methods). Overall 100 SNPs/MNPs/indels were called with a high confidence between the PAO1161 and the PAO1-UW reference. The remaining (heterozygous) variants were mostly SNPs (149/151) located in coding sequences of genes belonging to Pseudomonas Orthologous Groups (POG) possessing multiple members in the same strain (multiple orthologs within PAO1 genome)^6^. This group included 17 SNPs in *PA0723-PA0726* genes from the Pf1 cryptic prophage region syntenic to the genes within RGP42 element also present in PAO1161 genome^7^. Eight of these variants were also identified previously in MPAO1 and PAO1-DSM^7^, nevertheless these did not meet our quality criteria.

### Effect of SNPs, MNPs and indels

The high-confidence variants encompassed 52 SNPs, 6 MNPs, 15 deletions and 27 insertions. Of these 44 were mapping to the intergenic regions in PAO1-UW genome and 9 were synonymous (silent) mutations (Supplementary Table S1). Three mutations resulted in an introduction of premature stop codons (Table 1). Interestingly, these encompass *PA2735* gene, recently shown to encode a N6-adenosine DNA methyltransferase acting on a conserved sequence^32^, and *PA1939*, encoding a putative overcoming lysogenization defect (OLD) family nuclease containing an N-terminal ATPase domain and a C-terminal TOPRIM domain^33^. Since *P. aeruginosa* PAO1161 strain was selected as defective in its restriction system^11,17^, it is tempting to speculate that PA1939 could be the protein involved in degradation of the foreign DNA and together with PA2735 they could constitute the chromosomal restriction-modification system of *P. aeruginosa.*

**Table 1.** SNPs and indels identified in *P. aeruginosa* PAO1161 genome, resulting in expression of truncated proteins. The effect of a mutation is predicted using the PAO1-UW genome as a reference.

| Mutation effect | PAO1-UW position | Nucleotide change | AA change | length PAO1/PAO1161 | PAO1 gene | PAO1161 ID | Description |
| --- | --- | --- | --- | --- | --- | --- | --- |
| stop gained | 2121203 | C → T | W340* | 665 / 339 | PA1939 | D3C65_15950 | putative ATP-dependent endonuclease of the OLD family |
|  | 2356682 | CC → C | L173 | 182 / 172 | PA2141 | D3C65_14865 | CinA family protein |
|  | 3097884 | G → A | Q209* | 792 / 208 | PA2735 | D3C65_11725 | SAM-dependent DNA methyltransferase |
| frame-shift | 740419 | G → GC | V73A? | 381 / 124 | PA0683 (*hxcY*) | D3C65_22610 | putative type II secretion system protein |
|  | 1440623 | AA → A | K640 | 656 / 642 | PA1327 | D3C65_19175 | putative protease |
|  | 1835045 | G → GC<sup>†</sup> | S218S? | 249 / 226 | PA1685 (*masA*) | D3C65_17305 | enolase-phosphatase E-1 |
|  | 2807706 | CAGCCGGCC → C | aa1-78/35aa | 347 / 304 | PA2492 (*mexT*) | D3C65_13040 | transcriptional regulator |

<sup>†</sup> - SNP at this position in PAO1DSM / MPAO-1<sup>7</sup> but a nucleotide insertion in our study

Four of the identified sequence variants resulted in frame shifts, leading to expression of truncated proteins (Table 1). In comparison to PAO1, PAO1161 genome contains a frame shift in the *PA0683* (*hxcY*) gene encoding a component of the Hxc system, a type II secretion system active under phosphate-limiting growth conditions and dedicated to the secretion of low-molecular-weight alkaline phosphatases LapA and LapB^34–36^. The frameshift in PAO1161 genome results in expression of a protein with only 72 aa identical to PA0683 from PAO1-UW.

In contrast, indels identified in *PA1327, PA1685* and *PA2141* mapped to the 3’ part of the genes, resulting only in short sequence alteration of C-terminal regions of corresponding proteins, unlikely to affect their function (Table 1). This is highly plausible as the analysis of the protein sequence of PA1327, PA1685 or PA2141 orthologs (Pseudomonas Ortholog Groups: POG001263, POG003705, POG003333, respectively), showed that the PAO1161-like C-termini are typical for most orthologs from the corresponding groups.

The effect of 14 indels and 1 SNP is predicted as a shift in start or stop codon of the corresponding gene(s) leading to extension of the protein product in PAO1161 relative to PAO1-UW (Table 2). These also include changes in intergenic regions, like 413850 T → C mutation upstream of *PA0369*, modifying an in frame stop-codon and possibly leading to a translation of a longer protein. Similar, start codon changing role can be attributed to nucleotide changes within *PA2046-PA2047* and *PA4874-PA4875* intergenic regions, as well as to 5 other indels (Table 2). The effect of three PAO1161/PAO1-UW sequence differences is fusion of proteins encoded by adjacent genes in PAO1-UW genome into one protein in PAO1161. Additionally, the frame shift observed in PAO1-UW in *PA0748* gene, encoding an AraC type transcriptional regulator, is not observed in PAO1161. Notably, analysis of the sequences composing Pseudomonas Orthologs Group of corresponding PAO1 genes showed that in all cases the PAO1161-like protein products constitute a vast majority of all deposited sequences. One exception is the PAO1161 *relA* gene encoding a GTP pyrophosphokinase, in which a start codon shift was identified, compared to PAO1-UW (Table 2). RelA, together with SpoT, controls the level of guanosine 5’-triphosphate-3’-diphosphate (ppGpp) alarmone involved in bacterial adaptive responses and virulence^37^. Importantly, previous mass spectrometry analysis and iTRAQ quantification of PAO1161 proteome showed that RelA is produced in PAO1161 cells^38^.

**Table 2.** SNPs and indels identified in *P. aeruginosa* PAO1161 genome influencing the reading frames of genes. The effect of a mutation is predicted using the PAO1-UW genome as a reference. The sequences of PAO1161 proteins were compared with sequences from corresponding Pseudomonas Ortholog Group (POG) for PAO1 proteins.

| Position in PAO1-UW | Nucleotide change | PAO1 loci | PAO1161 loci | Effect of mutation |
| --- | --- | --- | --- | --- |
| 816531 | C → CC | PA0748 | D3C65_22250 | Corrects the reading frame of PA0748 (still frameshift orf). |
| 1215658 | G → GG | PA1122 | D3C65_20255 | Reading frame change after 125 aa. Fusion with 54 aa instead of 22 aa (PAO1). Similar to 498/504 ortologs from POG001082. |
| 2355771 | A → AG | PA2139 | D3C65_14875 | Reading frame change after 29 aa. Fusion with 42 aa instead of 11 aa (PAO1). Similar to 6/12 ortologs from POG008397. |
| 2753522 | G → GC | PA2451 | D3C65_13240 | Fusion of PA2452 and PA2451. No fusion only in PAO1-UW (POG003088). |
| 3083196 | A → AG | PA2727 | D3C65_11765 | Fusion of PA2727 with PA2728. Similar fusion observed for 366/372 proteins from POG002843. |
| 4539468 | G → GC | PA4058 | D3C65_04690 | Fusion of PA4059 with PA4060. No fusion only in PAO1-UW (POG001625). |
| 413850 | T → C | IG:<br>PA0369-PA0370 | D3C65_01955 | Removed TGA codon -12 to -10 from *PA0369* start codon. Predicted product has 30aa fused to N-terminus of PA0369. Similar to 297/303 ortologs from POG000360. |
| 2239555 | A → AG | IG:<br>PA2046-PA2047 | D3C65_15395 | Change in sequence preceding *PA2046*. Predicted product has 316 aa fused to N-terminus of PA2046. Similar to 289/298 orthologs from POG003399. |
| 5472415 | C → CG | IG:<br>PA4874-PA4875 | D3C65_26560 | Change in sequence preceding *PA4875*. Predicted product has 81 aa fused to N-terminus of PA4875. Similar to 292/299 orthologs from POG004503. |
| 1116213 | G → GC | PA1029 | D3C65_20745 | Removes start codon of *PA1029*. Predicted product has additional 24 aa fused to N- terminus of PA1029. Similar to 310/316 orthologs from POG000991. |
| 4888194 | A → AG | PA4360 | D3C65_23120 | Removes start codon of *PA4360*. Proposed product has additional 122 aa fused to N- terminus of PA4360. This extension is observed for most members of POG003953. |
| 1275766 | GA → G | PA1174 | D3C65_19970 | First 11 aa of PA1174 replaced by 16aa. Extended sequences in 343/350 ortologs from POG001132. |
| 3016844 | G → GC | PA2668 | D3C65_12075 | Reading frame and start codon changed. Predicted product has last 56 aa identical to C-terminus of PA2668. Extended sequences in 295/301 orthologs from POG008397. |
| 3919508 | G → GC | PA3503 | D3C65_07595 | Predicted product has last 210 aa identical to C-terminus of PA3503. Extended sequences in 69/73 orthologs from POG005142. |
| 1023047 | GGCAAGAT → G | PA0934<br>(*relA*) | D3C65_21265 | Start codon shift and M1VVKGR change. Only in PAO1161. |

Except nucleotide changes with a major effect of the corresponding protein products, numerous SNPs and indels resulting in amino acid substitutions or deletions relative to corresponding PAO1-UW proteins were identified (Table 3). One special case is PA2492 (*mexT*), where both a deletion and a SNP were observed in PAO1161 relative to PAO1-UW sequence (Table 1 and Table 3). MexT is a LysR type transcriptional regulator activating expression of the MexEF-OprN multidrug efflux system, extensively studied in the context of quorum sensing signalling and resistance to antimicrobial agents^39–42^. PAO1161 does not possess the 8-nt insertion within *mexT* (Table 1), observed in some PAO1 sublines^43^. Concomitantly, PAO1161 contains a SNP in the *mexT* gene, leading to F172I change which is also found in most of the strains with the active *mexT*^43^. A sequence variation was also observed in *mexS*, encoding a negative regulator of MexT^40,44,45^, resulting in D249N amino acid change, which was shown to not affect the protein function^40^.

**Table 3.**
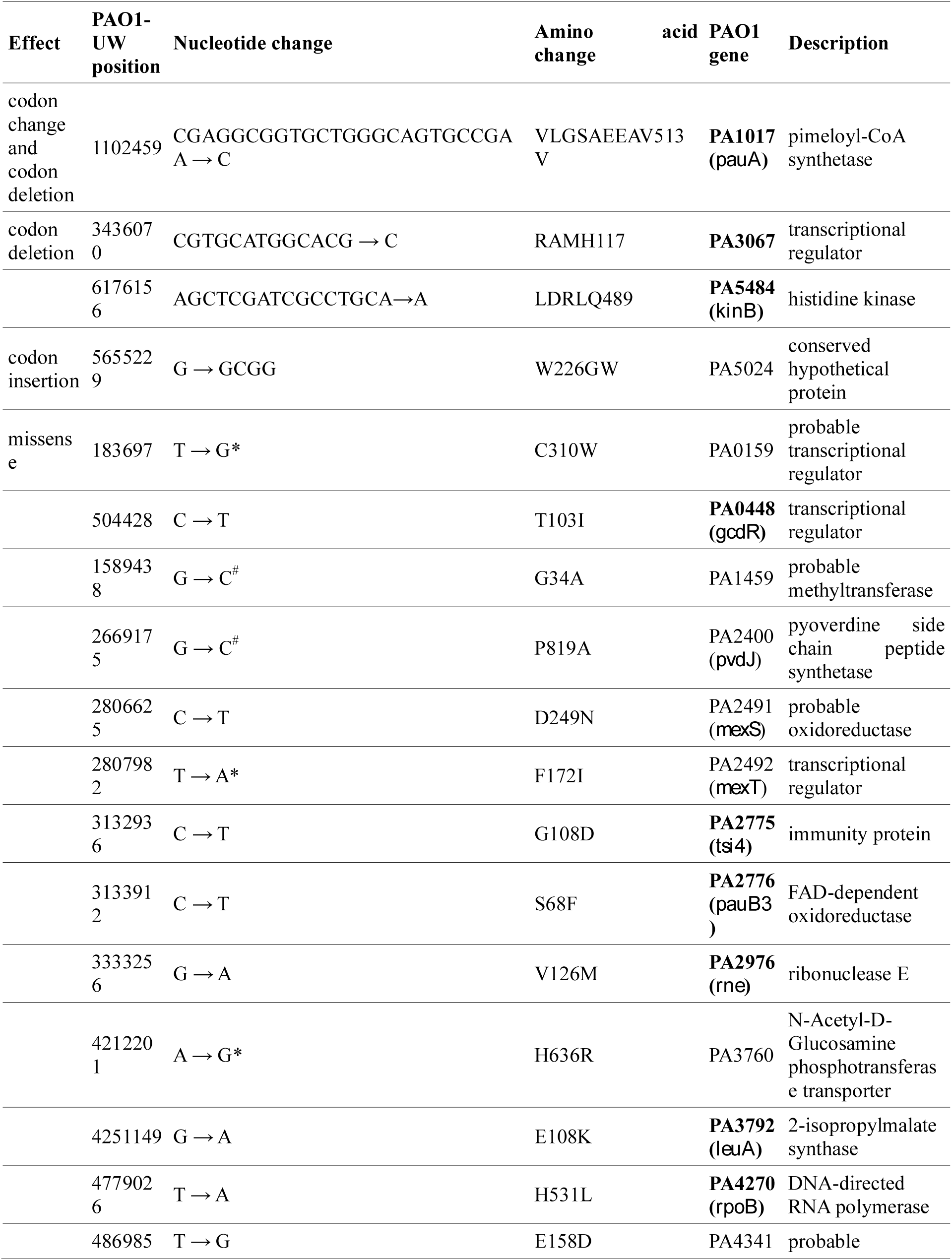
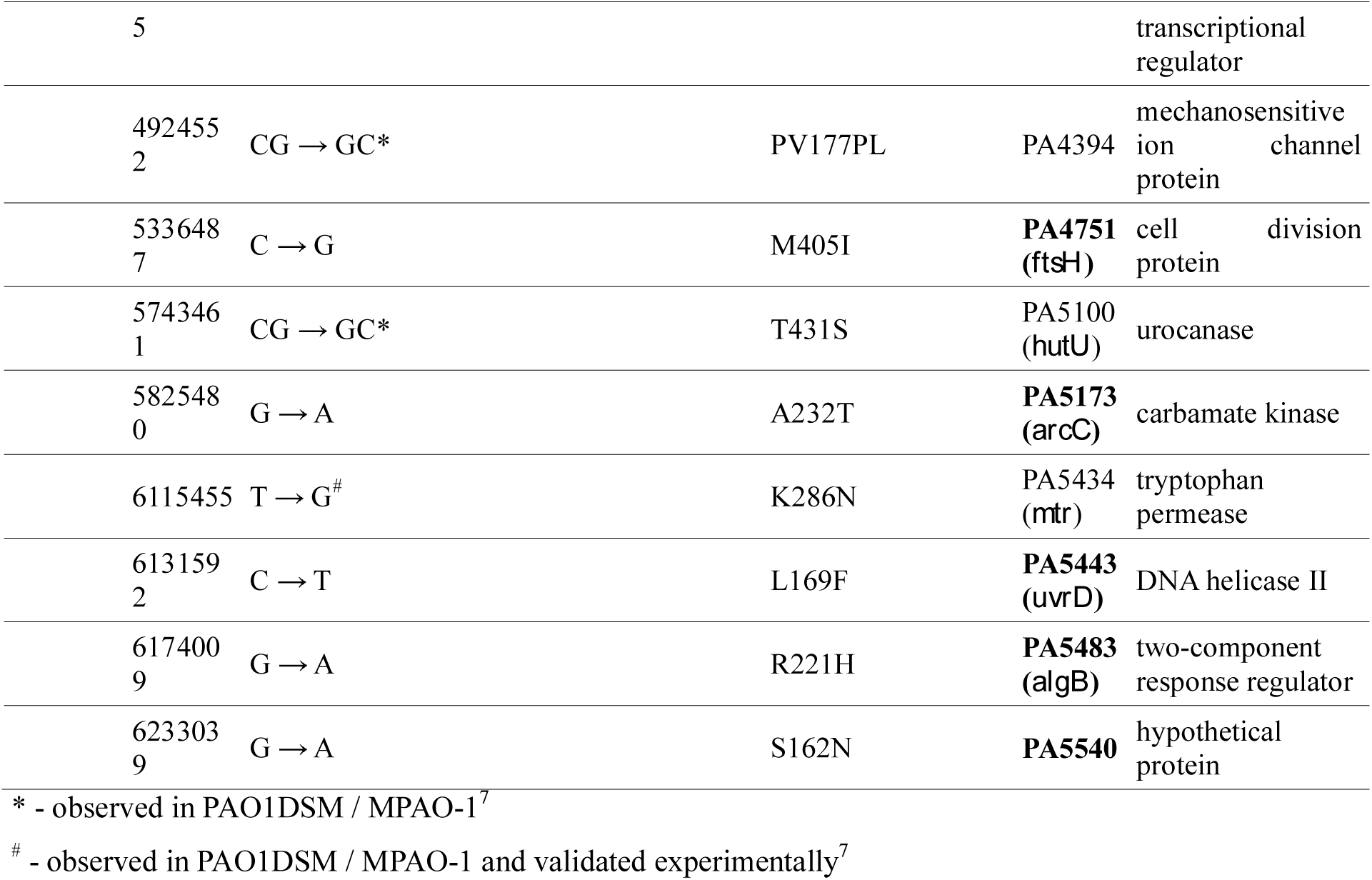
SNPs and indels identified in *P. aeruginosa* PAO1161 genome resulting in amino acid changes relative to corresponding PAO1-UW proteins. PAO1161 strain specific changes, identified by comparison of PAO1161 proteins with sequences in Pseudomonas Ortholog Groups encompassing corresponding PAO1 proteins, are indicated by bolded gene names.

PAO1161 strain derives from the strain PAO38 which was selected as the leucine auxotroph^46^ (Figure 1A). Genome sequencing of PAO1161 revealed that this strain possesses a mutation in *leuA*, encoding a putative 2-isopropylmalate synthase, resulting in E108K substitution. Analysis of Pseudomonas Ortholog Group of the *leuA* (POG001874) showed that, the only *P. aeruginosa* strains carrying this mutation are PAO579^46,47^ and PAO581^48^, two PAO38 derivatives (Figure 1A). To validate that this substitution causes the leucine auxotrophy, we exchanged PAO1161 *leuA* to the PAO1 allele. The allele replacement fully restored the ability of PAO1161 strain to grow on minimal medium without leucine (Figure 2), confirming that the E108K substitution in LeuA caused leucine auxotrophy.

**Figure 2.**
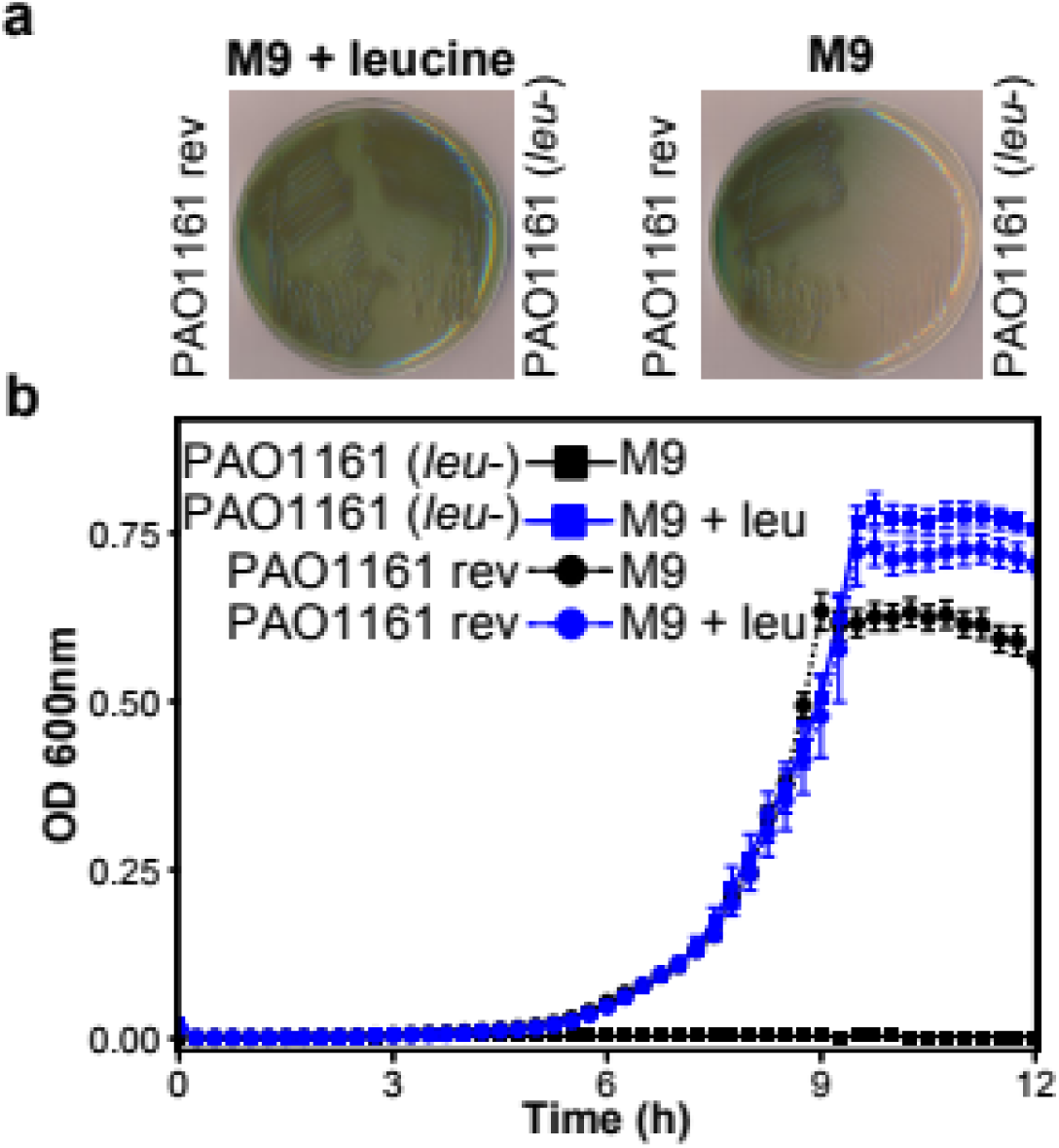
LeuA E108K substitution causes leucine auxotrophy in *P. aeruginosa*. PAO1161 *leuA* allele, carrying the mutation, was replaced with the PAO1 allele to yield strain PAO1161 rev (*leu*+). Growth of PAO1161 (*leu-)* and PAO1161 rev (*leu*+) strains on solid **(A)** and liquid **(B)** minimal medium containing 0.25% citrate with or without 10 μg ml^-1^ leucine. Data represent mean OD_600nm_±SD for 6 biological replicates.

*P. aeruginosa* PAO1161 used in this study was a rifampicin resistant clone^18^. Rifampicin binds to a conserved pocket on the β-subunit of RNA polymerase therefore blocking RNA transcript elongation^49^. Resistance to this drug results from mutations on the *rpoB* gene that change the structure of the pocket^49–51^. Our analysis revealed presence of a SNP in *rpoB*, encoding a DNA-directed RNA polymerase subunit beta, resulting in H531L substitution (Table 3). This amino acid change was frequently observed in spontaneous *P. aeruginosa* Rif^R^ mutants^52^, strongly indicating that this SNP confers PAO1161 strain with rifampicin resistance.

Interestingly, for 8 proteins the same changes were found in PAO1161 strain and in MPAO1 and / or PAO1-DSM^7^ relative to corresponding PAO1-UW proteins (Table 3). Fifteen changes seem to be PAO1161 strain specific (Table 3, bolded) as revealed by comparison of the sequences with other members from the corresponding Pseudomonas Ortholog Groups. These include codon deletions in ortologs of *pauA*, encoding a pimeloyl-CoA synthetase involved in biotin synthesis^53^ and *PA3067*, a MarR family transcriptional regulator. Amino acids deletion was also found in KinB, forming a two-component system with the transcriptional regulator AlgB involved in control of alginate and virulence factors production in *P. aeruginosa*^54–57^. The point mutation changing amino acid sequence R221H was also detected in PAO1161 *algB* (Table 3). It is tempting to speculate that the identified changes may attenuate the virulence properties of PAO1161 cells.

Other strain specific substitutions include T103I change in the transcriptional regulator GcdR regulating glutarate utilization and lysine catabolism^58^, A232T in the carbamate kinase ArcC involved in fermentative arginine degradation^59^, V126M in the ribonuclease E (*rne*) involved in mRNA turnover^60^. The G108D change was also identified in Tsi4 of PAO1161 relative to corresponding protein of PAO1-UW. This protein is a part of antibacterial effector-immunity pair together with type VI secretion exported toxin Tse4^61^. Overall, the identified substitutions could influence the structure and/or function of the corresponding proteins and should be considered in analyses involving these genes using PAO1161 strain.

### PAO1161 genome contains an ICE conferring resistance to mercury

PAO1161 was described as a strain containing the FP2 plasmid of IncP-8 incompatibility group, which confers the cells with mercury resistance^15^. Indeed, the strain used in our lab was exceptionally resistant to mercury, growing in LB medium supplemented with up to 200 μM HgCl_2_ (data not shown). Surprisingly, during the genome assembly no extra-chromosomal elements could be identified. Instead, an almost 108 kbp insertion in the chromosome, with a putative mercury resistance operon, was found (Figure 1B, Figure 3A). The insertion shows similarities (in sequence and organization/composition of operons flanking the putative integration site) to the PAPI-1 family of integrative conjugative elements (ICEs)^29,30,62^ abundant in *Pseudomonas* genomes. ICEs are mobile genetic elements, with a modular structure, encoding complete conjugation machinery (usually a type IV secretion system) allowing transfer of their genome to another host. They are reversibly integrated into a host genome and can be passively propagated during bacterial chromosome segregation and cell division^62–64^. PAPI-1 (108 kb, 115 orfs, integrated in *tRNA*_Lys_) was first described in the genome of highly virulent *P. aeruginosa* PA14 strain^65^.

**Figure 3.**
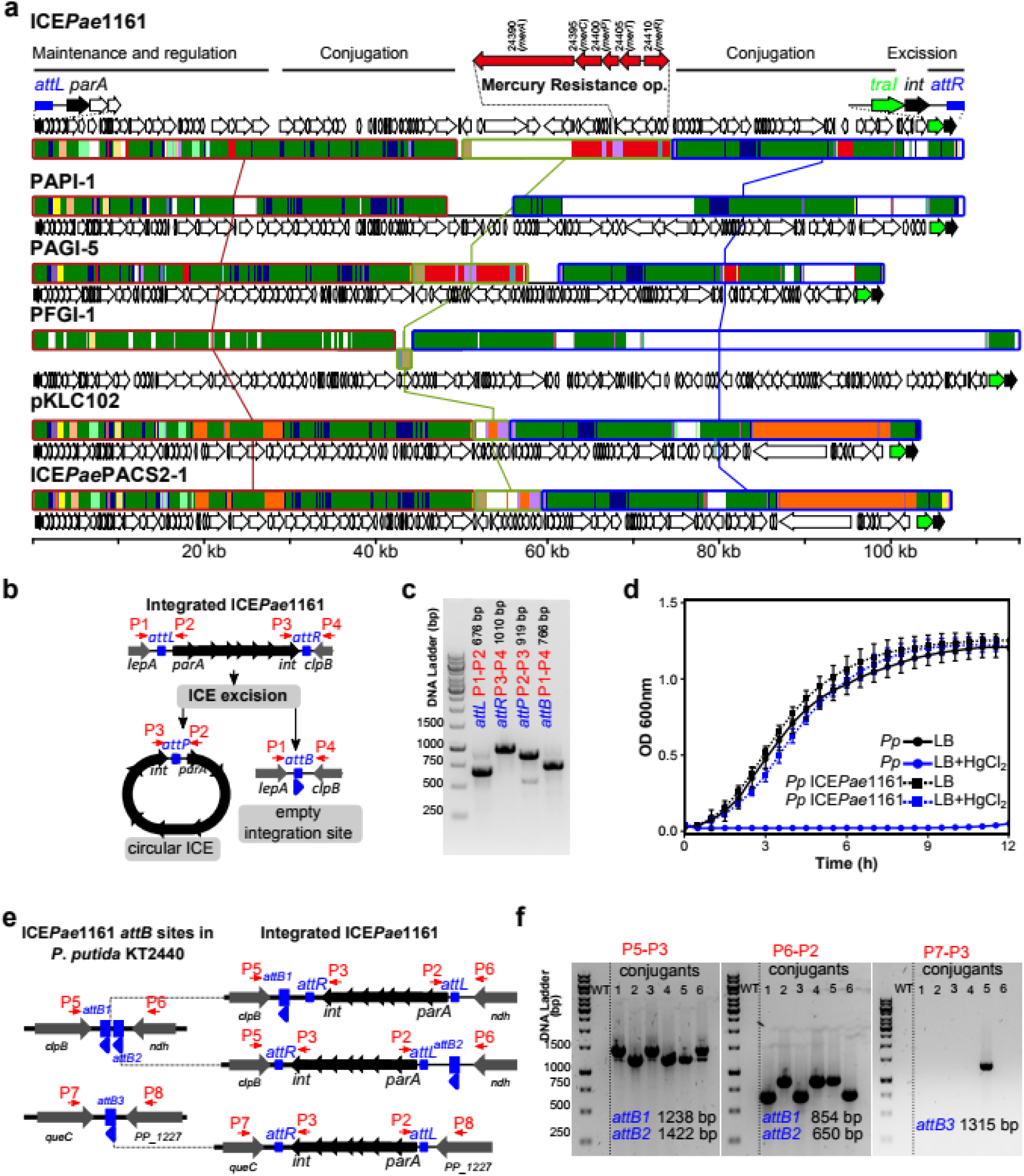
ICE*Pae*1161 identified in *P. aeruginosa* PAO1161 genome is a functional PAPI-1 family integrative and conjugative element conferring resistance to mercury. **(A)** Comparative genomics of ICE*Pae*1161 and selected PAPI-1 family ICEs. Mauve alignment of ICE*Pae*1161 and PAPI-1^65^, PAGI-5^81^, PFGI-1^82^, pKLC102^83^ and ICE*Pae*PACS2-1 (position 896693:1002644 of NZ_AAQW01000001)^29^ is presented. Three blocks of sequences which are free of genome rearrangements, such as inversions and duplications, are marked with rectangles connected with lines. Green segments indicate sequences conserved in all ICEs (backbone). Regions conserved among subsets of analysed ICEs are color coded. White region is specific only to one analysed ICE. Arrows indicate the location and orientation of coding sequences. Boundary genes and the mercury resistance operon are shown on top. **(B)** Schematic model of linear and excised (circular) ICE*Pae*1161. Primer binding sites are indicated in red. **(C)** PCR analysis of ICE*Pae*1161 excision. PCR was performed with indicated primer pairs flanking *att* sequences using PAO1161 genomic DNA as a template and products were separated on a 1.2% agarose gel. **(D)** Growth of *P. putida* KT2440 strain and transconjugants *P. putida* KT2440 ICE*Pae*1161-Sm^R^. Strains were grown in L broth with or without 40 μM HgCl_2_. Data represent mean OD_600nm_±SD for 6 independent transconjugants analysed in 3 biological replicates. **(E)** Genomic context of three potential ICE*Pae*1161 integration sites (*attB*) in *P. putida* KT2440 genome. The sites were identified based on the presence of 48 bp sequence flanking ICE*Pae*1161 in PAO1161 genome. Blue arrows indicate the orientation of the sequences. Schematic model of ICE*Pae*1161 integration at each *attB* site in orientation corresponding to the one observed in PAO1161 genome. **(F)** PCR analysis of ICE*Pae*1161 presence in *P. putida* KT2440. Genomic DNA of six independent transconjugants was used as a template in PCR using the indicated primers and products were separated on 1% agarose gels.

The element identified within PAO1161 genome, which we named ICE*Pae*1161, has an integration site within *tRNA*_Lys_ and PAPI-1 like organization of boundary operons: an operon starting with a gene encoding a putative ParA protein, at one end and an operon encoding a putative relaxase (TraI) and site-specific recombinase (Int) at the other (Figure 3A). Analysis of gene content (Supplementary Table S2), revealed that out of 120 predicted genes within ICE*Pae*1161, 102 genes are found in at least one other PAPI-1 like element, whereas orthologs of 41 genes are found in all ICEs analysed (Supplementary Table S2).

Integration of ICE into the chromosome as well as its excision is mediated by an ICE encoded site directed recombinase / integrase^66^. Recombination between an attachment site in the chromosome (*attB*) and the corresponding site on a circular ICE (*attP*) leads to integration of the element into the genome, now flanked by identical *attL* and *attR* sequences (Figure 3B). Excision of the ICE*Pae*1161 and the presence of a circular form was analysed using PCR with primers flanking the *att* sequences (Figure 3BC). The analysis confirmed occurrence of the circular ICE in PAO1161 cells (Figure 3C), indicating that the element can exist in two forms.

To verify the transferability of ICE*Pae*1161, we tagged it with a streptomycin resistance cassette (*aadA*). Subsequently, PAO1161 ICE-Sm^R^ strain was used as a donor in mating in static liquid cultures with *Pseudomonas putida* KT2440 as a recipient. The conjugants were selected on M9 plates supplemented with streptomycin, but lacking leucine to block the growth of donor cells. Streptomycin resistant *P. putida* clones were obtained with a low efficiency of 2 × 10^−7^ transconjugants / per donor cell.

To confirm that the *P. putida* conjugants show also enhanced resistance to mercury, attributed to the presence of *mer* operon within the ICE*Pae*1161, we analysed the growth of transconjugants in medium containing HgCl_2_. The recipient *P. putida* KT2440 cells were unable to grow at a HgCl_2_ concentration higher than 2 μM (data not shown). In contrast, the growth of streptomycin resistant *P. putida* transconjugants was not inhibited by the presence of 40 μM HgCl_2_ in the medium, confirming acquisition of mercury resistance (Figure 3D).

Finally, to confirm that the ICE*Pae*1161 integrated into *P. putida* KT2440 chromosome, we searched for putative *attB* sites in its genome using the 48 bp TGGTGGGTCGTGTAGGATTCGAACCTACGACCAATTGGTTAAAAGCCA sequence flanking ICE*Pae*1161 in PAO1161 chromosome. *P. putida* KT2440 genome contains three potential attachment sites designated *attB1*-*3*, two of them the *attB1* and *attB2* adjacent to each other (Figure 3E). A PCR analysis, using primer pairs specific to the ICE*Pae*1161/KT2440 chromosome junctions revealed predictably oriented ICE*Pae*1161 in the *P. putida* genome (Figure 3F). Among six individual transconjugants, the specific PCR products were observed preferentially for ICE integrated in one of adjacent sites, *attB1* and *attB2* with one clone also positive for ICE integration in *attB3* (Figure 3F). Overall, these data confirm the ability of the ICE*Pae*1161 identified in the PAO1161 genome to transfer to another host, integration into chromosome at a specific site and conferring mercury resistance.

### Conclusions

In this work we show that *P. aeruginosa* PAO1161 strain carries a PAPI-1 family integrative conjugative element capable of excision, transfer and integration in the genome of the related *Pseudomonas* strain. The ICE*Pae*1161 contains loci conferring mercury resistance, in the past attributed to the FP2 plasmid of IncP-8 incompatibility group.

The genome sequence of *P. aeruginosa* PAO1161 strain, a derivative of the reference PAO1 strain, was compared with the reference PAO1 sequence. The data indicate a number of sequence variants, likely affecting different aspects of cell physiology. Thus genome sequencing of laboratory strains is highly recommended to provide full insight of the genotype and help in the interpretation of phenotypes observed under the conditions studied.

## MATERIALS AND METHODS

### Strains and growth conditions

Bacterial strains, plasmids and oligonucleotides used in this work are listed in Supplementary Table S3. *P. aeruginosa* PAO1161 (*leu*^-^, *r*^-^, *m*^-^) was provided by B. M. Holloway (Monash University, Clayton, Victoria, Australia). *Escherichia coli* strain DH5α was used for plasmid manipulations and S17-1 was used to mate pAKE600^67^ derivatives into *P. aeruginosa*. Standard DNA manipulations were performed as described^68^. Templates for PCRs were prepared by boiling the cells pelleted from overnight cultures and resuspended in water.

Bacteria were grown in L broth^69^ at 37°C or on L agar (L broth with 1.5% w/v agar) supplemented with appropriate antibiotics. For selection of *E. coli* strains 150 μg ml^-1^ benzylpenicillin sodium salt in liquid medium, 300 μg ml^-1^ for solid media or 30 μg ml^-1^ streptomycin was used. *P. aeruginosa* and *P. putida* strains were selected by addition of 300 μg ml^-1^ carbenicillin, 300 μg ml^-1^ rifampicin or 150 μg ml^-1^ streptomycin to the medium. Growth analysis was performed in M9 minimal medium^68^ with 0.25% citrate or 0.1% glucose supplemented with 10 μg ml^-1^ leucine or 40 μM HgCl_2_ as indicated. Bacterial growth in 96-well plates was monitored by measurements of optical density at 600 nm (OD_600_) using a Varioskan Lux Multimode Microplate Reader and SkanIt RE 5.0 software (Thermo Fisher Scientific).

### Plasmids and strains construction

To construct prototrophic *P. aeruginosa* PAO1161 strain, which expresses LeuA without the E108K substitution, the suicide plasmid pKAB607 was constructed. Sequences flanking the region in PAO1161 chromosome were amplified using primer pairs 1#, 2# and 3#, 4#, respectively (Supplementary Table S3). Primers 2# and 3# contained sequence lacking the mutation observed in PAO1161 genome relative to PAO1-UW and introduced an AflII site, allowing selection of the allele. The obtained PCR fragments were digested with EcoRI, AflII and AflII, BamHI, respectively, and both fragments were ligated with EcoRI, BamHI digested pAKE600^67^ to yield pKAB607. *E. coli* S17-1 strain, carrying pKAB607 was used in the allele exchange procedure performed as described before^18^. Presence of modified allele was verified by AflII digestion of PCR amplified chromosomal *leuA*.

*P. aeruginosa* PAO1161 ICE-Sm^R^ strain was constructed by insertion of a streptomycin resistance cassette (*aadA*) between orfs D3C65_24365 and D3C65_24370 within ICE*Pae*1161 (Supplementary Table 2). To this end, first the fragments flanking the region of future insertion were amplified from PAO1161 genomic DNA using primer pairs 5#, 6# and 7#, 8#, respectively (Supplementary Table S3). The obtained PCR fragments were digested using MunI, HindIII and HindIII, BamHI, respectively, and the mixture was ligated with EcoRI, BamHI digested pAKE600 to yield pKAB608. The Sm^R^ cassette, excised with HindIII from pHP45O^70^, was ligated with HindIII digested pKAB608 to yield pKAB609. *E. coli* S17-1 pKAB609 strain was used as a donor in the allele exchange procedure with strain PAO1161 Rif^R18^. Integration of Sm^R^ cassette was confirmed by PCR.

### Genome sequencing

For genome assembly, reads obtained previously from sequencing of the input sample of *P. aeruginosa* PAO1161 strain from ChIP-seq experiments were used^19^. The sequencing yielded 23380926 reads (4676185200 nt of raw data). Obtained reads were quality-filtered using FastX toolkit (http://hannonlab.cshl.edu/fastx_toolkit/) and residual Illumina adapters were removed using Cutadapt (https://github.com/marcelm/cutadapt)^71^. A subsample (7000 000 reads) was used in draft genome assembly using Spades v3.11.1 (http://cab.spbu.ru/software/spades/) to estimate the size of PAO1161 genome.

Concomitantly, long reads were generated using MinION (Oxford Nanopore Technologies, Oxford, UK). Genomic DNA was sheared into 20 kb fragments using Covaris gTube (Covaris, Ltd., Brighton, United Kingdom) and the library was prepared using an ONT 1D ligation sequencing kit (SQK-LSK108) with the native barcoding expansion kit (EXP-NBD103). Nanopore sequencing was performed using the NC_48ⅅh_Sequencing_Run_FLO-MIN106_SQK-LSK108 protocol, R9.4.1 MinION flowcell and a MinION MkIB instrument. Raw nanopore data were basecalled using Albacore v2.3.1 (Oxford Nanopore Technologies, Oxford, UK). Reads were quality-filtered and Porechop (https://github.com/rrwick/Porechop) and NanoFilt^72^ were used for adapter removal. Overall, MinION sequencing yielded 161877 reads (2160720766 nt), with a median read length of 12489 nucleotides. Long nanopore reads were assembled in a hybrid mode with the Illumina data using Unicycler v.0.4.6^73^. Genome assembly resulted in 1 circular replicon of 6383803 bp. Errors identified in the assembly were verified by a PCR amplification of DNA fragments, followed by Sanger sequencing on an ABI3730xl Genetic Analyzer (Life Technologies, USA) using BigDye Terminator Mix v. 3.1 chemistry (Life Technologies, USA) followed by correction of the genome sequence using Seqman software (DNA Star, USA).

The assembled genome was annotated using the NCBI Prokaryotic Genome Annotation Pipeline^74^. The nucleotide sequence has been deposited in NCBI Nucleotide database (accession number CP032126).

### Genome analysis

Genome synteny between PAO1161 and PAO1-UW was visualized using EasyFig^75^. Structural variations, SNPs, insertions and deletion between the PAO1161 and PAO1-UW (NC_002516) sequences were identified using Nucdiff^31^. The effect of SNP/MNP and indels was predicted using snpEff^76^. Reads were mapped to the PAO1161 genome using Bowtie v2.3.4.2^77^ followed by a verification of the quality of the assembly in regions differentiating PAO1161 and PAO1-UW genomes. Percentage of a given base, relative to bases from all reads, at positions in PAO1161 genome corresponding to the identified SNPs, MNPs and short insertions was analysed using bam-readcount (https://github.com/genome/bam-readcount). Regions with short deletions were inspected using Integrative Genomics Viewer v2.4.9^78^. The variants present in more than 80% of the reads with an average mapping quality > 20, were considered as homozygous. In case of 86/100 variants the base frequency was higher than 99%.

### ICE analysis and conjugal transfer

Comparative analysis of ICEs and gene ortholog prediction was performed using Mauve v2015-02-13^79^. GenoplotR was used to visualize gene distribution^80^. ICE*Pae*1161 transfer from *P. aeruginosa* PAO1161 ICE*-*Sm^R^ to *P. putida* KT2440 was performed by growing the strains in L broth overnight at 37°C, harvesting the cells, resuspending in the same amount of medium and mixing in 1:2 (donor: recipient) ratio in plastic 1.5 ml tubes. The mixtures were incubated for 2 h at 37°C with shaking (300 rpm) and then 3 h without shaking, followed by centrifugation and resuspension of the cells in the same volume of 0.9% NaCl. To estimate the efficiency of ICE*Pae*1161 transfer a suspension was serially diluted in 0.9% NaCl and aliquots were spotted onto M9 minimal medium agar plates with 0.1% glucose and 150 μg ml^-1^ streptomycin without leucine. Lack of leucine allowed growth of *P. putida* KT2440 transconjugants and counter selected PAO1161 *leu*^*-*^ donor cells. Donor or recipient strain cultures treated in the same way were used to establish the titre of donor and recipient cells in the conjugation mixture. The transfer frequency was calculated as the number of transconjugants per donor cell.

## SUPPLEMENTARY MATERIALS

**Supplementary Table S1.** Synonymous and intergenic SNPs/MNPs and indels identified in PAO1161 / PAO1-UW genome comparison.

**Supplementary Table S2.** Gene content of ICE*Pae*1161. Orthologs from ofther ICE were predicted using Mauve^79^ assuming 70% of identity and at least 70% of the sequence coverage.

**Supplementary Table S3.** Strains, plasmids and primers used in this study.

## ACKNOWLEDGEMENTS

This work was funded by National Science Centre, Poland [2015/2015/18/E/NZ2/00675 granted to A.A.B. and 2013/11/B/NZ2/02555 granted to G.J.B.]. We thank Jan Gawor and Robert Gromadka (DNA Sequencing and Oligonucleotides Synthesis Laboratory, IBB PAS, Warsaw, Poland) for performing the DNA sequencing.

## AUTHOR CONTRIBUTIONS

A.A.B., A.K. and G.J.B conceived and designed the experiments. A.K., K.K., M.M and A.A.B. conducted the experiments. A.K., K.K., M.M., G.J.B and A.A.B. analyzed and interpreted the results. A.K. and A.A.B. performed bioinformatics analysis. A.K., G.J.B. and A.A.B. wrote the manuscript. All authors reviewed and approved the manuscript.

## COMPETING INTERSTS

The author(s) declare no competing interests.

